# Cumulative effect of aging and SARS-CoV2 infection on poor prognosis in the elderly: Insights from transcriptomic analysis of lung and blood

**DOI:** 10.1101/2020.06.15.151761

**Authors:** Upasana Bhattacharyya, B K Thelma

## Abstract

**Introduction:** The ongoing pandemic caused by severe acute respiratory syndrome coronavirus 2 (SARS-CoV- 2) has affected millions of people worldwide and with notable heterogeneity in its clinical presentation. Probability of contracting this highly contagious infection is similar across age groups but disease severity and fatality among aged patients with or without comorbidities are higher. We hypothesized that SARS-CoV-2 infection may augment aging-related gene expression alterations resulting in severe outcomes in elderly patients.

**Methodology:** We performed a comparative analysis of publicly available transcriptome data from Broncho Alveolar Lavage Fluid (BALF)/lung/blood of healthy aging group with i) COVID-19 patients; and ii) data of host genes interacting with SARS-CoV-2 proteins.

**Results:** We observed i) a significant overlap of gene expression profiles of patients’ BALF and blood with lung and blood of the healthy group respectively; ii) a more pronounced overlap in blood compared to lung; and iii) a similar overlap between host genes interacting with SARS-CoV-2 and aging blood transcriptome.

**Conclusions:** Pathway enrichment analysis of overlapping gene sets suggests that infection alters expression of genes already dysregulated in the elderly, which together may lead to poor prognosis. eQTLs in these genes may also confer poor outcome in young patients worsening with age and co-morbidities. Furthermore, the pronounced overlap observed in blood may explain clinical symptoms including blood clots, strokes, heart attack, multi-organ failure etc. in severe cases. This model based on a limited patient dataset seems robust and holds promise for testing larger tissue specific datasets from patients with varied severity and across populations.

## Introduction

In December 2019, severe acute respiratory syndrome coronavirus 2 (SARS-CoV-2), a novel coronavirus was first reported as a zoonotic pathogen, that causes COVID-19 in humans and is capable of human-to-human transmission^1^. Since then it has spread rapidly across the world and thus has been classified as a global pandemic by the World Health Organization^2^. As of June 10, 2020, there have been 7.04 million confirmed COVID-19 cases with 404000 deaths reported worldwide (https://ourworldindata.org) with continuing trend of sharp rise in both these categories in many countries/regions. The disease is highly heterogenous in its clinical presentation with the respiratory system being most commonly attacked and showing symptoms such as fever, cough, shortness of breath and fatigue. In addition, myalgia, neurological symptoms (headache, dizziness, and altered consciousness), ischaemic and haemorrhagic strokes, muscle injury and gastrointestinal symptoms without respiratory symptoms or fever are also reported in a subset of patients^3^. Furthermore, in COVID-19 patients other clinical complications such as hypercoagulable state with increased risk of venous thromboembolism^4^ and in some cases, a ‘cytokine storm’ comprised of tumor necrosis factor α (TNF-α), interleukins (IL) 6, 1β, 8,12, interferon-gamma inducible protein (IP10), macrophage inflammatory protein 1A (MIP1A), and monocyte chemoattractant protein 1 (MCP1)^5^ has been observed. Several clinical trials are ongoing but as of date, no drugs or other therapeutics have been approved by the U.S. Food and Drug Administration (FDA) to prevent or treat COVID-19^6^ and thus, treatment being provided is only empirical. On the other hand, clinical management includes infection prevention with control measures and supportive care including supplemental oxygen and mechanical ventilatory support when indicated^6^. Efforts to combat COVID-19 are severely hampered by grossly inadequate knowledge of several important aspects of the illness ranging from pathogen biology to host response, disease biology, target tissues and consequently treatment options. Therefore, there is an urgent need for a deeper understanding of the host–pathogen interaction biology of SARS-CoV-2, which in turn may offer important insights into general/personalized treatment strategies and management of the disease as well as development of new therapies.

Probability of contracting this highly contagious infection has been reported to be similar across age groups but the clinical manifestations in the elderly patients appear to be more severe^7–10^. Besides, increased mortality has been reported in COVID-19 patients with comorbidities such as cancer, hypertension, diabetes etc^10,11^. However, the reported global case-fatality rate (CFR) among aged patients with or without comorbidities are notably higher^7–10^. Collectively, these observations suggest that there may be a comparatively stronger association between age and poor prognosis of COVID-19, but this may well be multifactorial. Therefore, uncovering mechanism(s) underlying poor prognosis to SARS-CoV-2 infection among the affected elderly might be insightful for effective patient management and treatment. It is reasonably argued that with the aging process, elderly population may be susceptible to various disorders such as lung infection, pneumonia, cardiovascular disorder, cancer or general age-related health problems^12–15^. Several reports also suggest that dysregulation of cellular oxidant/antioxidant systems and higher baseline levels of proinflammatory mediators, such as C-reactive protein, TNF-α, IL−1β and IL−6 as well as decrease in several stem and progenitor cell populations (that provide the lung with a remarkable regenerative capacity upon injury) occur in elderly people that might overlap with viral mediated dysregulation^15–18^.

Based on this limited understanding, we hypothesize that expression of genes that changes during aging, might get further augmented on SARS-CoV-2 infection, leading to severe outcome in elderly patient. We attempted to explore this possibility by performing comparative transcriptomics using available data from two target tissues namely lung and blood in healthy aging group and COVID-19 patients. We also compared transcriptomic profile of aging lung and blood with host genes interacting with SARS-CoV-2 protein. We observed a significant overlap between gene expression profile in both lung and blood of healthy aging group and COVID-19 patients; which was much more pronounced in blood. Furthermore, there was a significant overlap between host genes interacting SARS-CoV-2 proteins in aging blood but not in lungs. These observations support previous reports that SARS-COV-2, primarily affects the respiratory system but its effects may manifest in blood leading to multiorgan failure in severe cases of COVID-19^5,19^.

## Methodology

### Hypothesis

A schematic representation of the hypothesis is shown below (Figure 1).

**Figure 1:**
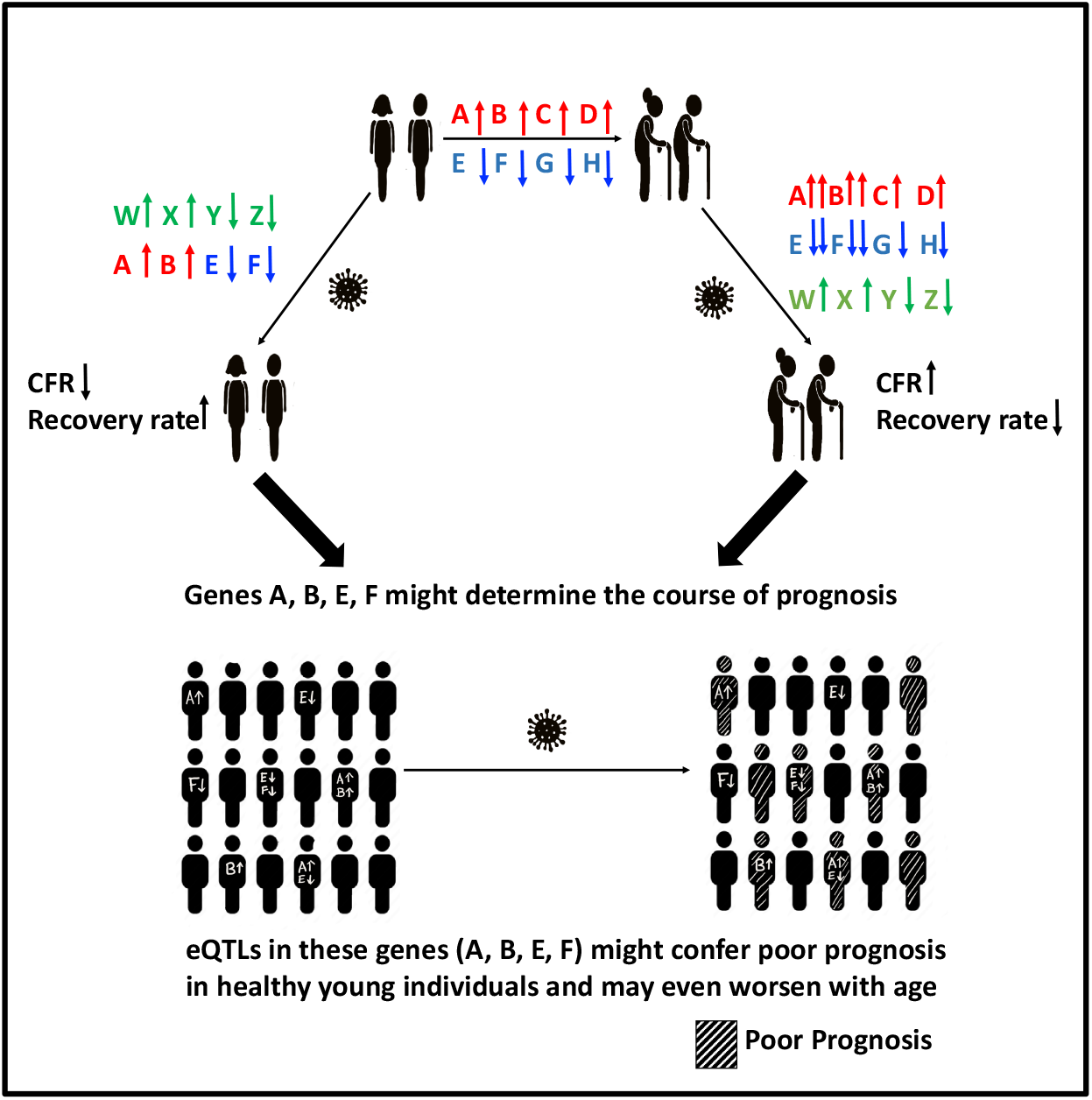
A schematic view of the hypothesis of differential gene expression for poor prognosis in the elderly COVID-19 patients

## Study Design

### Identification of candidate genes

In order to identify genes that determine poor prognosis in the elderly COVID-19 patients, we performed a comparative analysis of gene expression data collected from Broncho Alveolar Lavage Fluid (BALF)/lung/blood from COVID-19 patients and healthy aging group; and data of host genes interacting with SARS-COV-2 proteins with transcription profile of aging lung and blood (resources are mentioned below). Overlaps between the following groups were documented: i) Patients’ BALF with healthy aging lung; ii) Patients’ PBMCs with healthy aging blood; iii) Host genes interacting with viral proteins with healthy aging lung and blood.

### Statistical analysis

Statistical significance of these overlaps, if any, were tested using hypergeometric test (http://nemates.org/MA/progs/overlap_stats_prog.html). Basic equation to find probability of finding an overlap of genes using the above-mentioned program is provided in supplementary text.

### Pathway analysis

Pathway enrichment of the significantly overlapping genes identified above, was done using EnrichR^20,21^ which is an integrative web-based and mobile software application that currently includes 180184 annotated gene sets from 102 gene set libraries and various interactive visualization approaches to display enrichment results using the JavaScrit library, Data Driven Documents.

### eQTL analysis

eQTL variants of genes from significantly overlapping gene-sets for respective tissues (lung and blood) were obtained from GTEx version 8 (https://gtexportal.org). With a view to obtain a universally applicable biomarker, a comparable minor allele frequency of the markers would be ideal and such a suitability was tested using FST or fixation index. This was done using 1000 genome phase 1 data with the help of an online tool SPSmart (http://spsmart.cesga.es) with FST i) 0 to 0.05 representing low; (ii) 0.05 to 0.15 - moderate; iii) 0.15 to 0.25 – high; and >0.25 - very high, genetic differentiation.

### Identification of druggable targets

Finally, these genes identified above were screened for their druggability with FDA approved drugs using DGIdb website (http://dgidb.org).

## Resources

### A. Age associated genes

i. Differentially expressed genes (DEGs) in lung and blood tissue identified by a recently published study using RNA-Seq based transcriptome profiles from human donors of various ages from GTEx^22^ were obtained.
ii. DEGs from two stage transcriptomic study performed in blood, based on meta-analysis data from six different studies (*n*=7,074 samples) in the discovery phase and 7,909 additional whole-blood samples in the replication phase^23^ were obtained.

These two datasets for blood transcriptomics were merged and all the protein coding genes (and not any small RNAs such as miRNA, lncRNA etc) that are reported in either study were considered. All genes showing a different direction of expression change were removed from the analysis. A total of 2877 and 2283 protein coding genes were found to be up and down regulated respectively in blood from healthy aging group. Similarly, a total of 363 and 592 protein coding genes were found be up and down regulated respectively in aging lung. These genes have been subsequently referred to as “age-associated DEGs”.

### B.1. SARS-CoV-2 associated genes

DEGs in COVID-19 positive patients, henceforth referred to as “SARS-CoV2-associated DEGs” were obtained from two recent studies:

i. DEGs in BALFs identified comparing laboratory-confirmed COVID-19 patients (SARS2) (n = 8, median age 50.5 years) with healthy controls without known respiratory diseases (healthy) (n = 20)^24^.
ii. DEGs in PBMCs and BALF identified by comparing three COVID-19 patients (median age 37 years) and three healthy donors^25^.

BALF transcriptomics from these two datasets for were merged and all the protein coding genes (and not any small RNAs such miRNA, lncRNA etc) that are reported in either study were considered. All genes showing a different direction of expression change were removed from the analysis.

### B.2. Host genes interacting with SARS-CoV-2 proteins

Host genes that were found to be interacting with SARS-COV-2 viral proteins henceforth referred as “SARS-CoV2-interacting genes” were collected from recent studies mentioned below.

i. 332 high-confidence SARS-CoV-2-human protein-protein interactions (PPIs) were obtained by analyzing the data generated by expressing 26 of the 29 SARS-CoV-2 proteins in HEK293 cells in a recent study^26^.
ii. Computation based interactome data consisting of 125 proteins (94 host proteins) and 200 unique interactions which were generated using available sequences for viral protein candidates (wS, wORF3a, wE, wM, wORF6, wORF7a, wORF7b, wORF8, wN, and wORF10) as well as literature based mining^27^.

## Results

A comparative analysis of DEGs in the different sample sets described under study design (Figure 2), revealed significant overlaps between them (Table 1, Supplementary Table 1) and are briefly presented below. The most notable findings include i) significant overlap (p<1.4E-04) between the up regulated SARS-CoV-2 associated DEGs in patients’ BALF and up regulated age associated DEGs in healthy aging; ii) significant overlap (p<6.53E-07) between the up regulated SARS-CoV-2 associated DEGs in patients’ PBMCs and up regulated age associated DEGs in healthy aging blood; iii) nominally significant overlap (p<0.03) between the down regulated SARS-CoV-2 associated DEGs in patients’ PBMCs and down regulated age associated DEGs in healthy aging blood and iv) significant overlap between the SARS-CoV-2 interacting genes and up (p<0.002)/down (p<1.04E-06) regulated age associated DEGs in healthy aging blood (Table 1, Supplementary Table 1).

**Figure 2:**
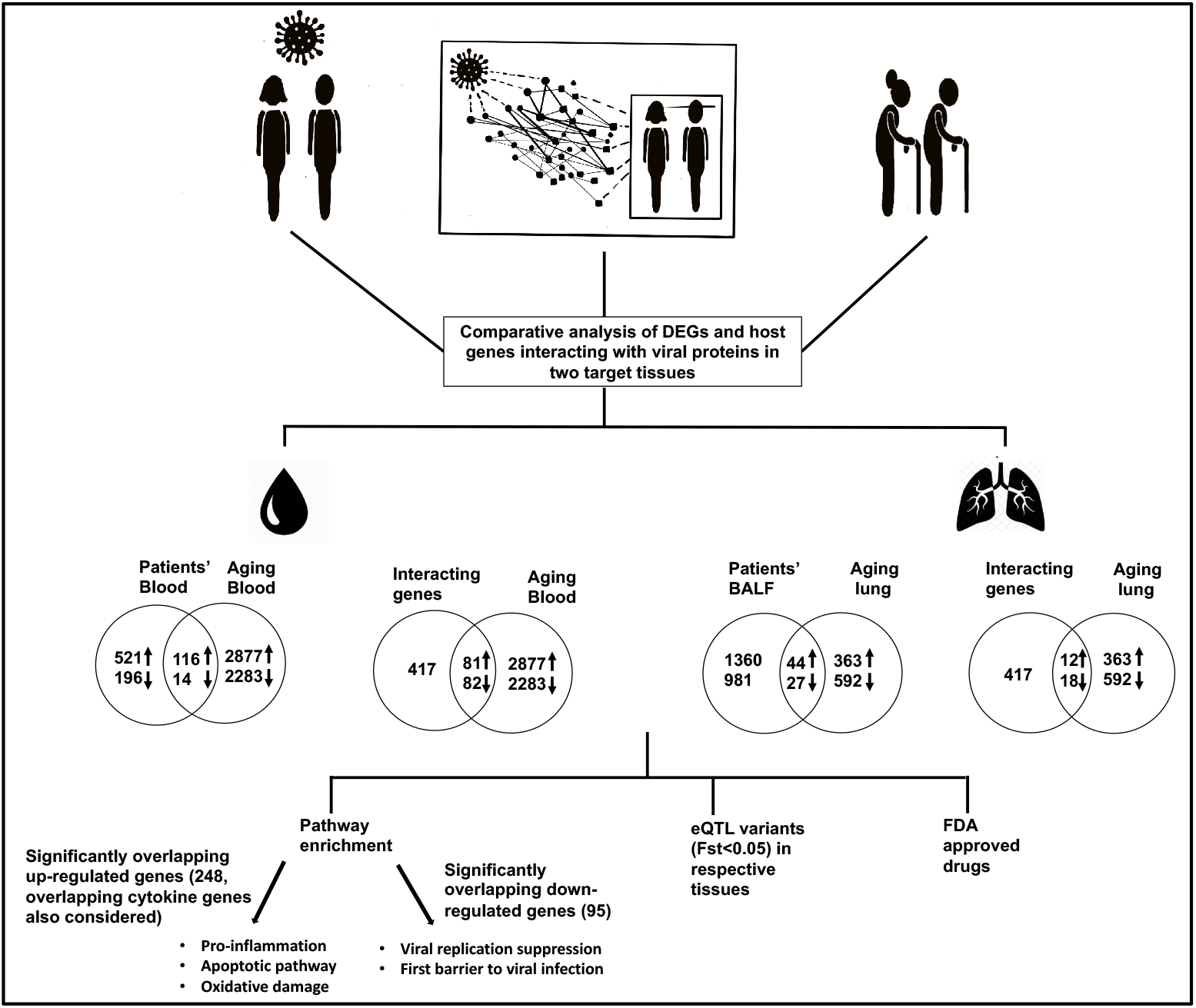
Shows the workflow and results of the comparative transcriptomics across different study groups

**Table 1:**
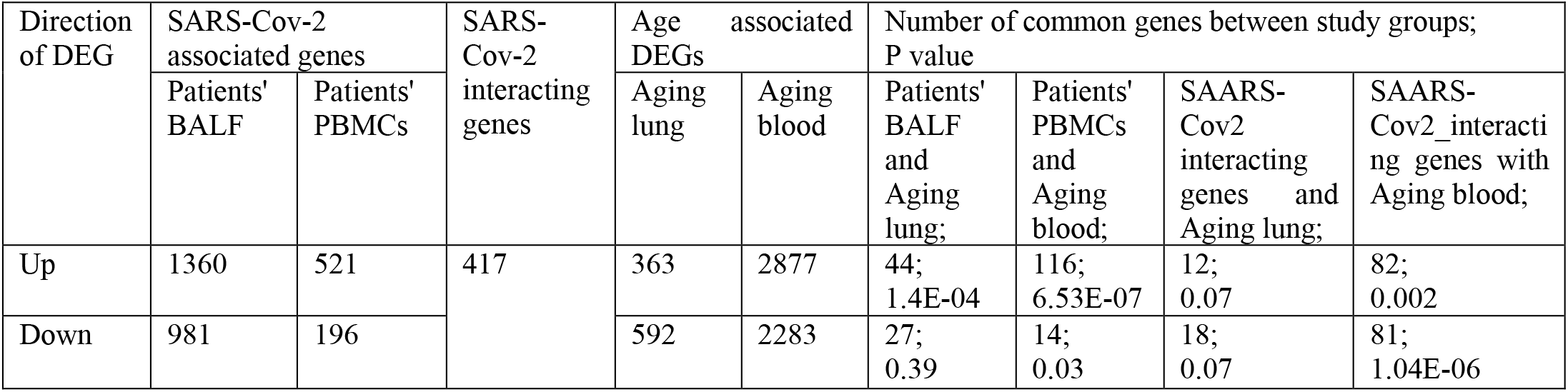
shows the results of the comparative analysis of differentially expressed genes (DEGs) across different sample sets

| Direction of DEG | SARS-Cov-2 associated genes |  | SARS-Cov-2 interacting genes | Age associated DEGs |  | Number of common genes between study groups; P value |  |  |  |
| --- | --- | --- | --- | --- | --- | --- | --- | --- | --- |
|  | Patients' BALF | Patients' PBMCs |  | Aging lung | Aging blood | Patients' BALF and Aging lung; | Patients' PBMCs and Aging blood; | SAARS-Cov2 interacting genes and Aging lung; | SAARS-Cov2_interacting genes with Aging blood; |
| Up | 1360 | 521 | 417 | 363 | 2877 | 44;<br>1.4E-04 | 116;<br>6.53E-07 | 12;<br>0.07 | 82;<br>0.002 |
| Down | 981 | 196 |  | 592 | 2283 | 27;<br>0.39 | 14;<br>0.03 | 18;<br>0.07 | 81;<br>1.04E-06 |

### Pathway enrichment

As the number of genes in each gene-set was very small to identify the enriched pathways, if any, all the genes from significant gene-sets and showing change in same direction were considered together for pathway enrichment analysis. Cytokine genes that are frequently found to be upregulated in patients and which overlapped with healthy aging expression profiles were also included. Six of the 21 up-regulated cytokine genes in patients, overlapped with aging related genes in the blood but none with aging lung. Up regulated genes in the healthy aging group that overlap with SARS CoV-2 associated genes and SARS-CoV-2 interacting genes were seen to be enriched in a range of signaling pathways including p53, chemokine and cytokine mediated inflammation, EGF receptor, TGF-beta, AGE-RAGE, Toll-like receptor mediated, NF-kappa B, VEGFA-VEGFR2 and genes involved in ROS in triggering vascular inflammation, oxidoreductive damage, T cell polarization, lung fibrosis, Chronic Obstructive Pulmonary Disorder (COPD), local acute inflammatory response (representative pictures at Figure 3; Supplementary Table 2). Down regulated genes in the healthy aging group that overlap SARS CoV-2 associated and SARS-CoV-2 interacting genes were seen to be enriched in pathways such as PI3K-Akt-mTOR-signaling, membrane trafficking, HIV and influenza RNA transport, ISG15 antiviral mechanism, cellular export machinery that interacts with NEP/NS2 (representative pictures at Figure 4; Supplementary Table 3).

**Figure 3:**
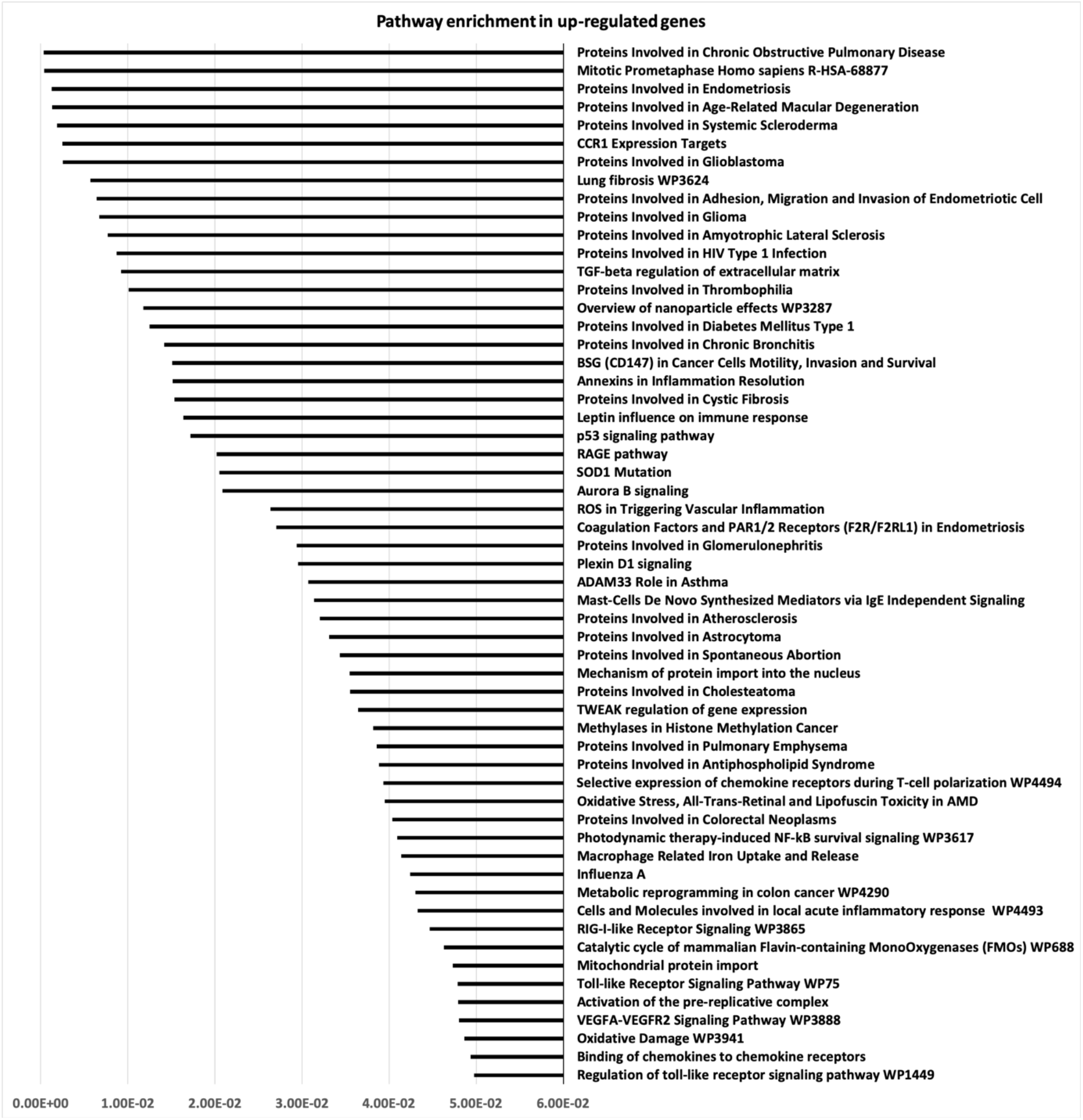
Shows result of pathways enrichment analysis of up regulated genes in the healthy aging group overlapping with SARS CoV-2 associated genes and SARS-CoV-2 interacting genes

**Figure 4:**
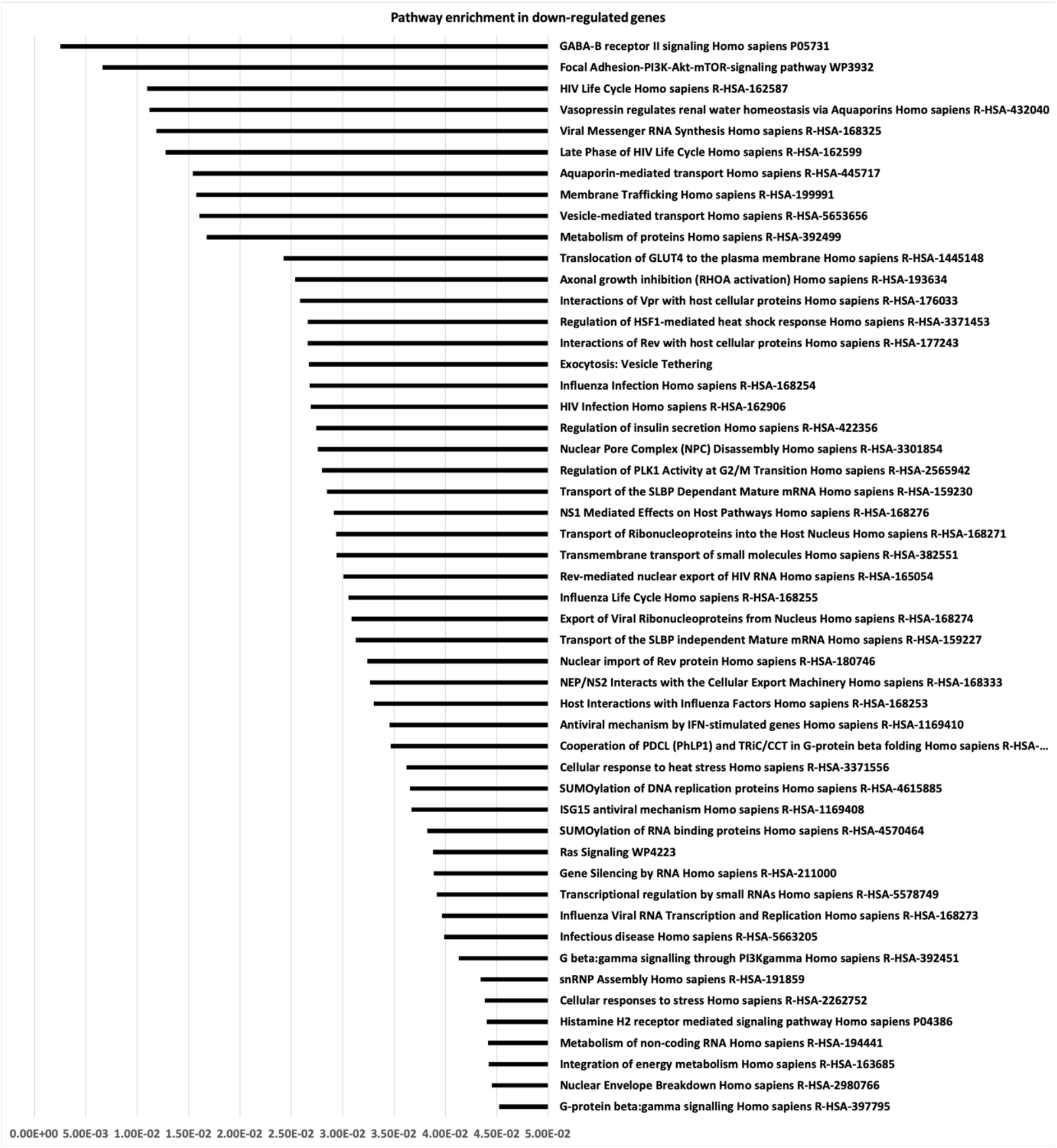
Shows result of pathways enrichment analysis of down regulated genes in the healthy aging group overlapping with SARS CoV-2 associated genes and SARS-CoV-2 interacting genes

### eQTL analysis

eQTL variants in genes from significantly overlapping gene-sets from the respective tissue namely lung and blood were identified (Table 2, Supplementary Table 4). Fst test were performed to identify variants with low genetic differentiation among different population. A large number of eQTL variants with Fst<0.05 were found in each group (Table 2, Supplementary Table 4). Further analysis of these may enable identification of variant(s) which could be used as a biomarker(s).

**Table 2:** shows number of genes with significant eQTL variants in each group and number of variants with Fst<0.05 in them

| Direction of expression change in patients and healthy aging group | Up-regulated |  |  |  | Down-regulated |  |
| --- | --- | --- | --- | --- | --- | --- |
| Gene-set common between study groups (n) | Patients' BALF and aging lung (44) | Patients' PBMCs and aging blood (116) | SARS-CoV-2 interacting genes with aging blood (82) | Cytokine and aging blood (6) | Patients' PBMCs and aging blood (14) | SARS-CoV-2 interacting with aging blood (81) |
| Number of eGenes in GTEx dataset (n) | 22 | 58 | 44 | 3 | 8 | 49 |
| Total number of eQTL variants | 1163 | 2717 | 2118 | 72 | 90 | 2269 |
| Number of variants with $F_{st} < 0.05$ | 395 | 1215 | 1611 | 40 | 34 | 1302 |

### Identification of druggable gene targets

It may be mentioned that SARS-CoV-2 interacting genes were excluded from this analysis since they have already been reported previously^26^. Efforts to identify novel druggable targets for FDA approved drugs from among 259 genes from significantly overlapping gene sets between different groups shown above (Table 1), yielded a total of 48 druggable genes mostly from the immune system related pathways. A total of 205 FDA approved drugs could potentially target them (Supplementary Table 5).

## Discussion

COVID-19, the recent pandemic has affected individuals of all age groups as well as all ethnicities. However, poor prognosis has been witnessed in the elderly effected group with or without comorbidities^7–10^. The overall CFR for cases with age 70 to 79 years is 8.0% and for cases with age 80 years and above it is 14.8%; which is strikingly higher compared to 0.4% in cases below 50 years of age, based on a study that included 72314 cases from China^28^. It is well known that with aging there is a notable increase in circulating pro-inflammatory cytokines even in the absence of an immunological threat and also a reduction in proteins that maintain homeostasis of the immune system, thus entailing a greater risk for many diseases (cancer, cardiovascular and neurodegenerative disorders, COPD, lung cancer, interstitial lung disease etc)^14,15,29,30^. Thus, we reasoned that comparing naturally occurring, age-dependent transcriptional changes with those observed in COVID19 patients and/or known SARS-COV-2 interacting proteins may provide insights into the age-associated poor prognosis in COVID-19.

Three noteworthy findings emerged from our study: i) a significant overlap was witnessed between DEGs in patients’ BALF/blood and in aging lung and blood; ii) this overlap was more pronounced in blood compared to lung; and iii) a similar overlap between SARS-CoV-2 interacting proteins and DEGs in aging blood but not in lung (Table 1) and warranting discussion. These overlaps suggest a cumulative effect of aging and COVID-19 infection on expression of important genes which may be conferring poor prognosis in the elderly. Results of pathway enrichment analysis in both up-regulated and down-regulated genes (Figures 3,4; Supplementary Tables 2,3) lend good support to this interpretation. We observed that genes involved in pathways such as pro-inflammatory, apoptotic, T cell polarization were up-regulated and those involved in viral replication suppression or the first barrier to viral infection remained downregulated in the elderly (Figures 3, 4; Supplementary Tables 2,3). Based on this, it may be proposed that on SARS-CoV-2 infection, expression of these genes changes further, thus causing lowered immunity, enhanced pro-inflammatory response and apoptosis which in turn may lead to poor prognosis.

As for the second novel observation, the higher extent of overlap between SARS-CoV-2 associated/SARS-CoV-2 interacting genes and DEGs in blood among the heathy aging group, may explain the range of clinical symptoms including high prevalence of blood clots, strokes and heart attack as well as multi-organ failure in a subset of severe patients reported early on during this pandemic^3,4^. Our observations are corroborated by a recent report of SARS-CoV-2 mediated damage to endothelial cells lining the blood vessels probably leading to blood clotting, strokes and heart attacks^19^. Furthermore, our observations suggest an over activation of T cell polarization and TGF-beta pathways in affected elderly patients (Figure 3), which can cause functional exhaustion of T cells^31,32^. A recent study suggests T cell counts are reduced significantly in COVID-19 patients, and the surviving T cells appear functionally exhausted in severe cases^33^.

Taken together these observations, eQTLs in the genes identified in our study (Table 2, Supplementary Table 4) may also confer poor prognosis in young patients but this may worsen with age and co-morbidities. However, their utility as prognostic biomarkers across different populations may be assessed only when more patient data become available and these genes are characterized in COVID-19 patients. On the other hand, variants with Fst>0.05 may be tested for correlation with population specific severity. Additionally, identification of genes involved in influenza and HIV infection in our study (Figure 3,4; Supplementary Table 2,3) may be of considerable relevance for drug repurposing for treatment of COVID-19 patients. In summary, our analysis i) identified probable candidate genes for poor prognosis among the affected elderly group; ii) identified potential druggable targets as well as FDA approved drugs opening the possibility of drug repurposing; and iii) seems to provide early explanation for manifestation of blood related symptoms and probably multiorgan damage. Finally, though these leads are preliminary based on a very limited patient dataset, the model holds promise to be tested as and when more tissue specific patient derived data from different age groups and/or from different populations become available.

Limitation: Our study is limited by non-availability of large tissue specific transcriptomic datasets of COVID19 patients across age groups, and with varying severity and from different populations to validate the hypothesis.

